# Generic workflow for a rapid and easy design of strain-specific PCR and qPCR primers, applied to the assessment of bacterial strains survival in soil

**DOI:** 10.1101/2024.05.06.592701

**Authors:** A. Polrot, J. Béguet, M. Devers-Lamrani, F. Martin-Laurent, A. Spor

**Affiliations:** Agroécologie, INRAE, Institut Agro, Univ. Bourgogne, Univ. Bourgogne Franche-Comté, Dijon, France

**Keywords:** Primer design, strain specificity, qPCR, full workflow, user-friendly

## Abstract

**Background:** Detecting bacteria at the strain level is crucial in microbiology. Although qPCR is widely used, designing strain-specific primers remains a challenge due to nucleotide sequence similarities among related strains.

**Methods and Results:** This paper introduces a simplified, web-based workflow for designing strain-specific primers using publicly available microbial genomes. The method does not require advanced bioinformatics skills and can be applied using a basic computer. Primers designed using this workflow are applied to assess the survival of two close *Bacillus* strains in soil microcosms.

**Conclusion:** The workflow offers an accessible solution for accurate bacterial strain detection, and fills a gap for researchers without specialized training in bioinformatics.

## Introduction

Detecting and identifying specific bacterial strains within different microbiota is crucial in clinical, applied and environmental microbiology. A wide range of scientific applications – disease diagnostic, environmental monitoring, food safety, bioremediation, biocontrol studies – requires specifically detecting microbes at the strain level. Over time, a large variety of techniques has been developed for the specific detection of microbes, ranging from traditional culture-based methods to advanced molecular techniques [1-4]. In this respect, techniques like quantitative PCR (qPCR) have become increasingly popular [5, 6] because they rapidly and sensitively detect specific DNA sequences associated with microbial strains. Primer design is a critical step in the development of a new qPCR assay. However, designing primers specific to a particular bacterial strain remains a significant challenge because nucleotide sequence similarities among closely related strains can result in non-specific amplification and overestimated abundance of the targeted strain. At the era of high-throughput sequencing, more and more full microbial genomes are being sequenced and made publicly available. Their increased accessibility facilitates the design of specific primers by relying on unique sequences specific for a given strain rather than focusing on dissimilarities in more conserved gene regions. Nonetheless, a remaining challenge in the design of strain-specific primer pairs is the need for expertise in bioinformatics, which can be a significant barrier for researchers who lack this skillset. Existing methods for designing primers rely on complex algorithms and software, which can be difficult to use without specialized training.

This paper aims to fill this gap by providing an easy workflow to design strain-specific primers from genomes, by exclusively using web-based software. This workflow does not require advanced skills in bioinformatics, but only a broad understanding of FASTA files handling and BLAST. It can be performed on any basic computer connected to the internet, using web-based interfaces. To validate the utility of this workflow, we have designed primers for four bacterial strains with a shared function— degradation of atrazine—and for two strains within the same species of *Bacillus*. These last primers were further validated for qPCR, and their application was demonstrated in an experiment assessing the survival of the two *Bacillus* strains in soil microcosms.

## Materials and methods

### *In-silico* primer design workflow

The workflow is provided in figure 1. The first step consists in extracting sequences specific to the strain of interest from the existing bacterial genomes *via* a galaxy public server [7]. Galaxy is a web-based analysis platform freely accessible via public servers. These servers can simply be accessed through a web browser and present a user-friendly interface in which the tools can be selected and their parameters easily modified. The Galaxy Europe server was used for this study, but any of the three principal UseGalaxy public instances can be used (Galaxy Main, Galaxy Au, Galaxy EU), among others.

**Figure 1:**
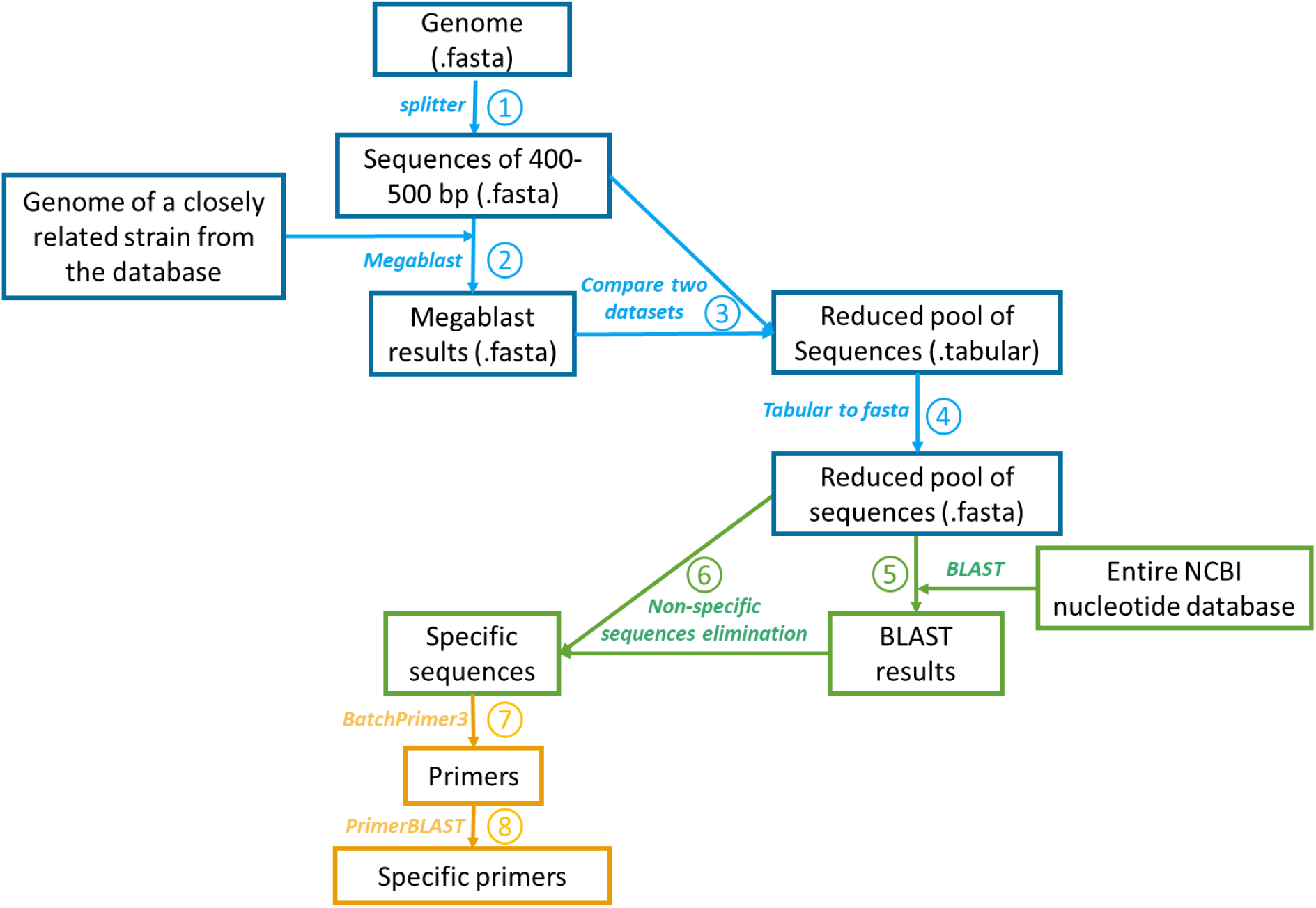
Workflow for the design of specific primers from bacterial genomes using web-based software and tools. Blue, steps performed using galaxy; green, steps performed using BLAST ; yellow, actual primer design steps. The complete description of the parameters is given in supplementary file 1.

The genomes are first cut into 500 base-pair (bp) pieces using the “splitter” tool. Details of the parameters for each Galaxy tool are provided in supplementary file 1. If no suitable primer is found in the later stages, the split size can be modified: decreasing it may help retrieve more specific sequences previously eliminated because of adjacent non-specific locations; increasing it will modify the locations of the cuts and may reveal more suitable primer locations. Next, the output of the splitter tool is matched against the genome of a closely related strain using “megablast” from the “NCBI BLAST + BLASTN” tool. The “closely related strain” is chosen among the available strains of the same species in the database, and a blast can be run on a random portion of the genome of interest to select the first matching strain. The sequences resulting from the megablast search are removed from the split genome using “Compare two Datasets”. This step results in a set of sequences that are unique to the genome of interest compared to the genomes of the closely related strains. Then, the data is transformed in FASTA format using the “Tabular-to-FASTA” tool and downloaded. These remaining sequences are finally matched against the entire NCBI nucleotide database using the BLAST web interface, and only the sequences without any match to nucleotides in the database are kept. The sequences resulting from the previous steps are specific to the strain of interest and can be used as a reservoir to design primer pairs specific for unique sequences without a match.

For the actual primer design, Batchprimer3 web-based software is used with the parameters described in supplementary file 1 [8]. If no candidate primer is found, some parameters can be adjusted depending on the desired application for the primers. For example, if the intended use is PCR and not qPCR, the parameters can be loosened: product size can be increased to several hundreds of bp, primer size can go from 18 to 24 bp, and primer Tm can range from 55 to 60 °C (making sure that the pair have a Tm within 3 °C of each other). Finally, each resulting primer pair is checked using the primer-BLAST tool of NCBI and the parameters described in supplementary file 1 [9].

### *In vitro* primer screening

Various soil DNA samples were used as PCR matrices to test the specificity of all the primers (Supplementary files 3 and 4). For *Bacillus* strains A and B, the primers giving the neatest amplification of DNA from the matching strain and no amplification at all of DNA from the non-matching strain and from various soil samples (Supplementary file 4) were retained. qPCR standards were made and tested in the qPCR assay, as described in Supplementary files 5 and 6. Limits of quantification (LOQ) were calculated using dilutions of DNA extracts of each strain in ten replicates. The retained LOQ was the last dilution that gave a variation coefficient lower than 30%.

### Final primer validation

For the final validation of the primers specific to *Bacillus* strains A and B, three agricultural soils with different physico-chemical properties were collected in France (Supplementary file 7). The non-sterile soils were inoculated with 10^8^, 10^6^ or 10^4^ cells *per* gram of dry soil (cells/g soil dw) in four replicates. Non-inoculated negative controls were also prepared. DNA was directly extracted using the DNeasy 96 PowerSoil Pro Kit (Qiagen). Quantifications and dilutions to 0.5 ng/µL were performed, as well as an inhibition test. qPCR assays targeting specific sequences were carried out to detect and quantify the abundances of the two strains.

### Application to a survival trial

Soil microcosms were set up for the evaluation of the survival of *Bacillus* strains A and B. 50 grams (dry weight) of three agricultural soils with different physicochemical properties were weighed in glass microcosms. To prepare the inoculation of the microcosms, each strain was cultivated in LB growth medium in 2 L conical flasks, agitated at 120 rpm for 24 hours at 28°C. The bacterial culture was then rinsed with 0.9% NaCl and adjusted at concentrations suitable to inoculate the microcosms at two doses: a high dose of 1 x 10^8^ Bacteria/g of soil dw and a low dose of 5x10^5^ bacteria /g of soil dw. The four treatments (two strains, two doses) and a control without inoculation were set up in four replicates. All the microcosms were then adjusted at 50% of water holding capacity and incubated at 20°C. Four sacrificial sampling were done at time = 0 days, time = 14 days, time = 28 days and time = 56 days. DNA was then extracted, diluted and quantified by qPCR as described in the previous paragraph.

## Results and discussion

### Primers design, screening and LOQ determination

The workflow described in this study is inspired from [10] and [11]. This latter study fully describes a workflow for the design of specific primers using user-friendly tools based on the analyses of open reading frames (ORFs), which limits the number of unique sequences to coding regions of the genome. Non-coding regions represent only a small proportion of the genomes of prokaryotes. However, their genomes are by nature less conserved because they are not subjected to the same selection pressure as coding DNA: they are selected for their size rather than for their function [12]. Therefore, non-coding regions may be more likely to harbor strain-specific sequences. In addition, the workflow described in [11] involves the downloading of the entire NCBI database, which can take several hours or finally fail.

Our workflow provides a solution to avoid the handling of such a big file (> 50 Go) and is inspired from another study which suggests to first compare the genome of the bacterial strain of interest with the genomes of related strains available in the databases so as to eliminate the major part of its genome shared with other genomes [10]. However, these authors used in-house scripts and non-user-friendly software, making the method difficult to reproduce by unexperienced users.

At first, the workflow described here was also applied to design primers specific to bacterial strains sharing the same function. These strains capable of atrazine degradation (*Pseudomonas* sp. strain ADPe, *Arthrobacter* sp. strain TES, *Variovorax* sp. strain 38R, and *Chelatobacter* sp. strain SR38) have been described, and their genome is available in the NCBI database [13, 14].

Totals of 292, 18, 620 and 517 primer pairs were found for *Pseudomonas* sp. strain ADPe, *Arthrobacter* sp. strain TES, *Variovorax* sp. strain 38R, and *Chelatobacter* sp. strain SR38, respectively. For all of them, the first primer pair tested (described in Supplementary file 2) efficiently discriminated them. PCR assays with these primer sets did not yield amplicons from various environmental soil samples (Supplementary file 3), showing sufficient specificity to track each strain in the samples, e.g. in the context of biodegradation experiments.

To go further, the workflow was used to design specific primers for two closely related isolates of *Bacillus sp*. (strains A and B, 98.54% similarity, 84% alignment length coverage). The primers for these two strains were also validated by qPCR.

Ten primer pairs were obtained for strain A, and six for strain B. Each pair targeted different unique sequences of non-coding DNA, highlighting the advantage of the workflow. Only one pair *per* strain was retained after the PCR screenings (described in Supplementary file 2).

The LOQ were determined at 3 target sequence copies *per* reaction and 240 target sequence copies *per* reaction for strains A and B, respectively, i.e. 3 and 240 target sequence copies *per* ng of DNA, respectively, if 1 ng of DNA is used as a template for the qPCR.

### Primers validation for soil detection

The results of the final validation of the specific primers by qPCR detection of the strains in soil samples are showed in Figure 2. The two strains were successfully detected in soil microcosms inoculated at 10^6^ and 10^8^ cells/g soil dw. No specific amplicon was detected in the non-inoculated control soil microcosms or in those inoculated at 10^4^ cells/g soil dw. The qPCR quantifications of the two strains were about 10 times higher than those resulting from plate count of the bacterial culture used to inoculate the soils. This can be explained by the high efficiency of the primers but also the presence of dead cells or chains in the plates that led to underestimated counts. For the quantification of strain B, more DNA was introduced in the qPCR reactions to bring the copy number above the quantification limit. The statistical analyses, including the Kruskal-Wallis test, revealed no significant difference in the values for strain A (p = 0.98) or B (p = 0.66) among the different soils tested, suggesting the absence of a significant soil effect on the detection of these two strains under our experimental conditions.

**Figure 2:**
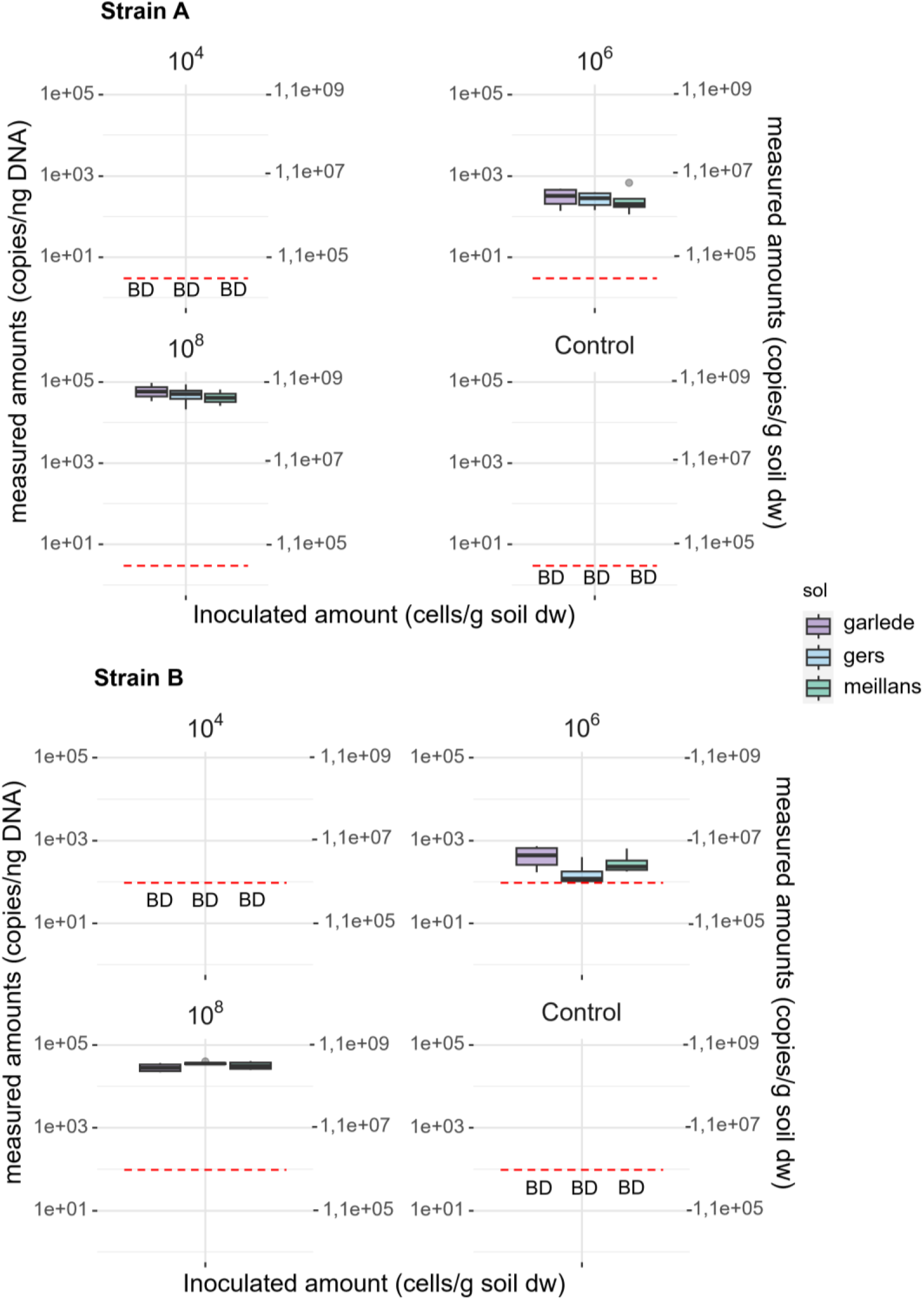
Boxplot of the measured copies of specific sequences per gram of soil in three soils for three inoculation values of our two strains of interest (A and B). Each boxplot represents four replicates. Dotted line, limit of quantification; BD, values below the limit of quantification.

The primer design method used here was successful to set up qPCR assays in order to detect and quantify two strains of *Bacillus* in DNA extracts from three soils using strain-specific primer pairs. Neither strain was detected in the soil microcosms inoculated at 10^4^ cells/g soil dw. This quantification is constrained by the DNA extraction and dilutions steps. DNA extracted from soil usually contains high quantities of PCR inhibitors, so that DNA has to be diluted before processing [15]. Lower detection limits in soil can be achieved using DNA extraction kits that are more efficient to remove soil inhibitors or do not require high dilution levels before qPCR processing [16]. Nonetheless, the quantification limits measured in copies *per* ng of DNA or copies *per* reaction is in accordance with those commonly found in the literature for qPCR primers [17].

### Application of strain specific primers to evaluate two *Bacillus* strains survival in soil microcosms

The results of the evaluation of *Bacillus* strains A and B survival in soil microcosms are shown in figure 3. The two strains were successfully detected for the high and low doses at the beginning of the experiment, however, at the low dose the values fell under the limit of detection from day 14 and remained undetectable until the end of the experiment. For the high dose, after a quick drop of two logs, the strains seemed to stabilize around 5x10^5^ copies/g of soil dw. Note that for strain B, the values of the low dose at 0 days and the values for the high dose from time 14 days until the end of the experiment are under the limit of quantification. The presence of specific amplification attests the presence of the strain, but the quantification values cannot be interpreted. For strain A, on day 56, quantification was challenging due to values being too close to the detection limit, despite successful specific amplification. This experiment further underlines the limitations of qPCR technique when working on environmental samples such as soil, due to the high quantification limits linked to the qPCR method applied to soil DNA extracts. Although we managed to gain some insights onto the survival capacities of strain A and B inoculated at 10^8^ bacteria per gram of soil dw, their fate is unclear at a lower dose of application, which is more representative of practical application.

**Figure 3:**
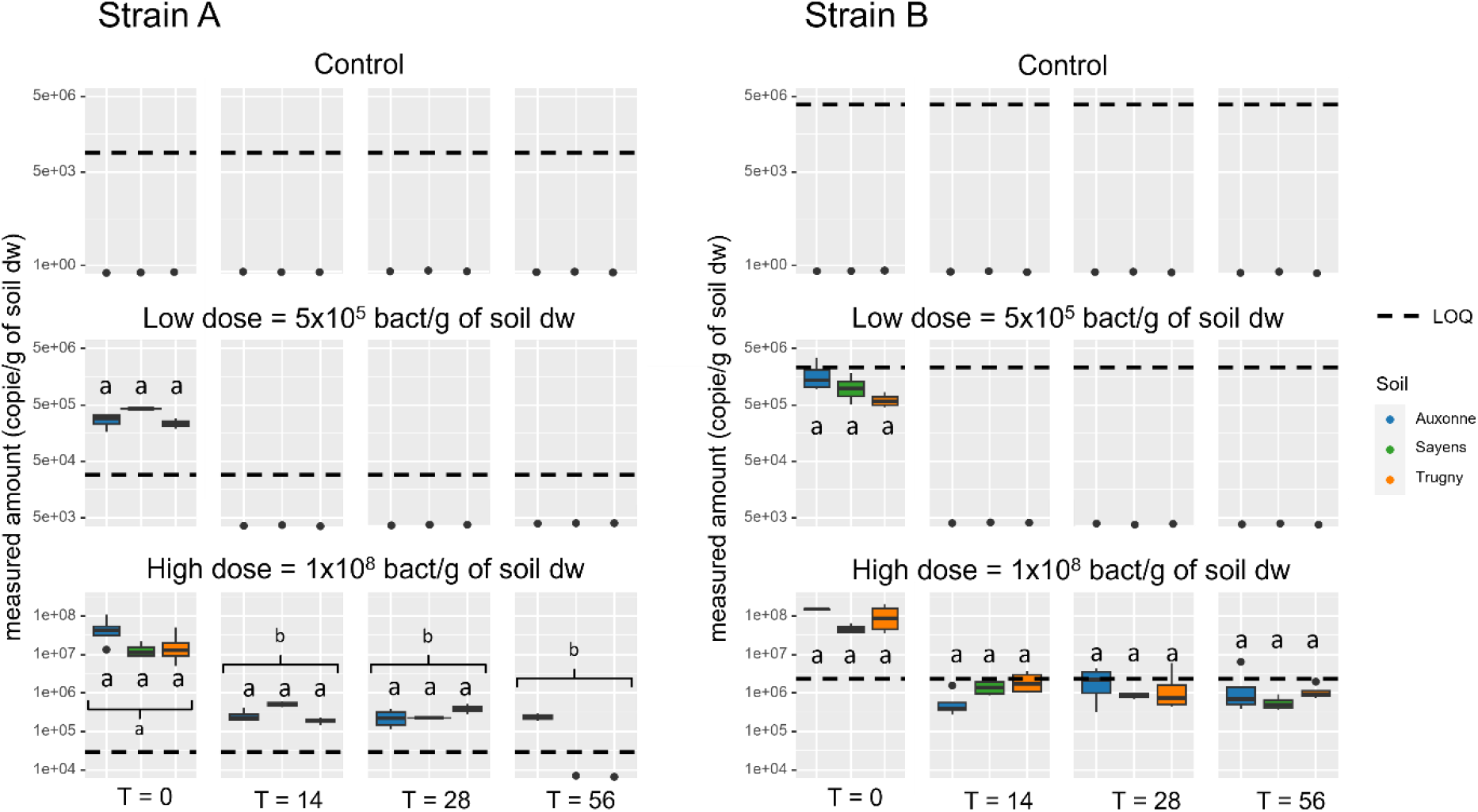
Survival potential of Bacillus strain A and B measured by qPCR using strain specific primers. Each boxplot represents four replicates. Each column represents a sampling time and each line a treatment. Letters show the statistical differences under the test of Kruskal-Wallis (alpha = 0.05).

The literature shows contrasting results regarding the survival of bacterial strains added to soil. While some studies indicate a rapid decline in population, reaching the limit of detection within less than 4 weeks [18, 19], the majority report survival for at least 2 months [20–25], but usually record a decline below the limit of detection over prolonged experimental durations [24, 25]. Nevertheless, certain studies have documented stabilization of bacterial counts even after extended periods, such as 100 days for a strain of *Listeria* [26] or 120 days for a strain of Burkholderia [27]. The survival of introduced bacteria in soil is influenced by a complex interplay of factors including soil physicochemical properties, application dosage, and the structure and diversity of indigenous microbial communities [28, 29]. In our study, conducting a longer experiment would be necessary to ascertain the fate of the two Bacillus strains after 2 months. However, the high dosage of application and their ability to sporulate could potentially account for their stabilization until that point.

## Conclusion

Overall, the described easy-to-go workflow is effective to rapidly and easily design strain specific primers. These primers can then be used to detect and quantify a strain of interest from environmental microbiota with the appropriate qPCR assays. Despite certain methodological limitations, particularly at lower inoculum levels, this workflow was successfully applied to advance our understanding of microbial survival in soil environments and sets a foundation for future applications in environmental microbiology.

## Statements and Declarations

### Data Availability Statement

The authors declare that the data supporting the findings of this study are available within the paper and its Supplementary data file. Should any raw data files be needed in another format they are available from the corresponding author upon reasonable request.

### Author Contributions

Conceptualization: AP, MD, FML, AS; Formal analysis: AP; Investigation: AP, JB; Supervision: MD, FML, AS; Writing – Original Draft: AP; Writing – Review & Editing: FML, AS, JB, MD.

