## Supplementary Data for "Generic workflow for a rapid and easy design of strain-specific PCR and qPCR primers, applied to the assessment of bacterial strains survival in soil"

Supplementary file 1: Strain specific primer design workflow

1) Create a galaxy account, Galaxy Europe was used in the present study but another region can be chosen depending on study location (https://usegalaxy.eu/)
2) Upload your genome and the genome of a closely related strain (in .fasta format) from databases by clicking “Upload Data” and dropping the files in the window than opens up, confirm the uploading process by clicking “start”.

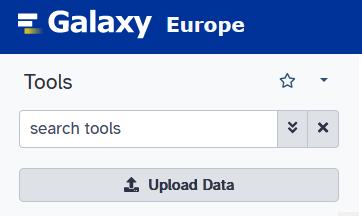

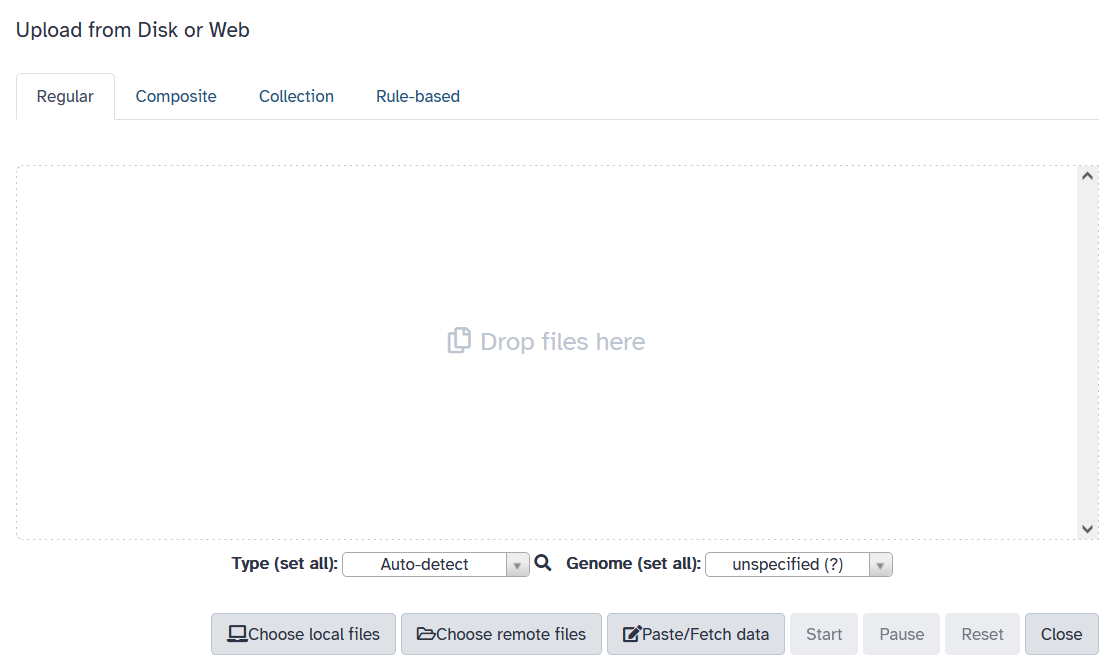

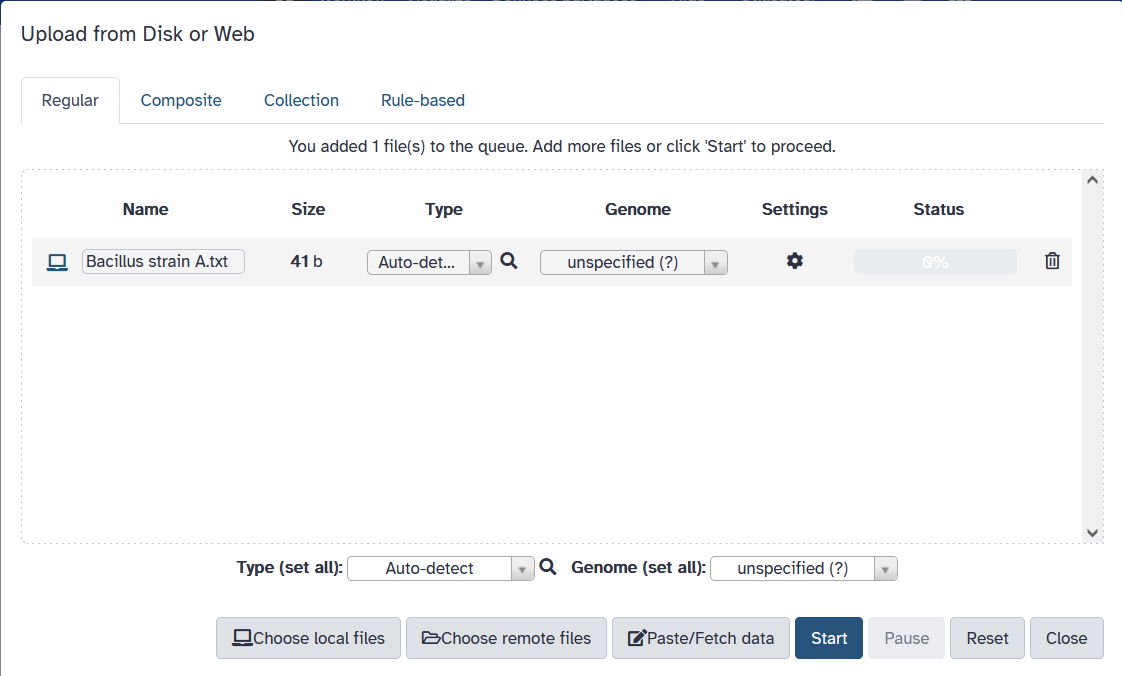

3) Search for the “splitter” tool using “search tool” and run it with the below parameters, the size to split at can be adjusted.

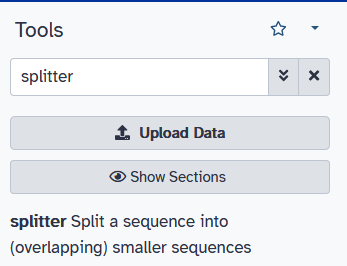

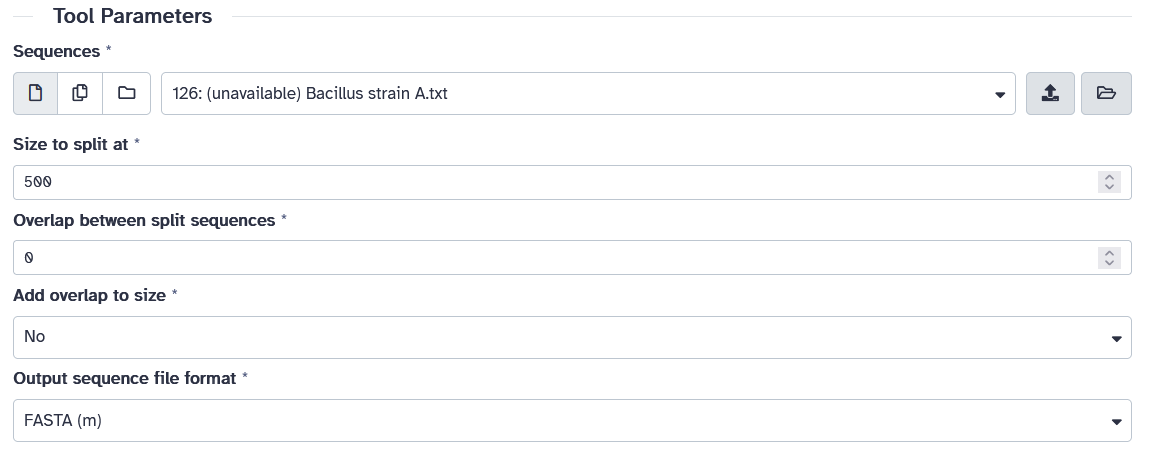

4) Search for the “NCBI BLAST+ blastn” tool and run it using the data generated by the splitter tool and a genome of a closely related strain from the strain you are designing the primers from. Use the below parameters.

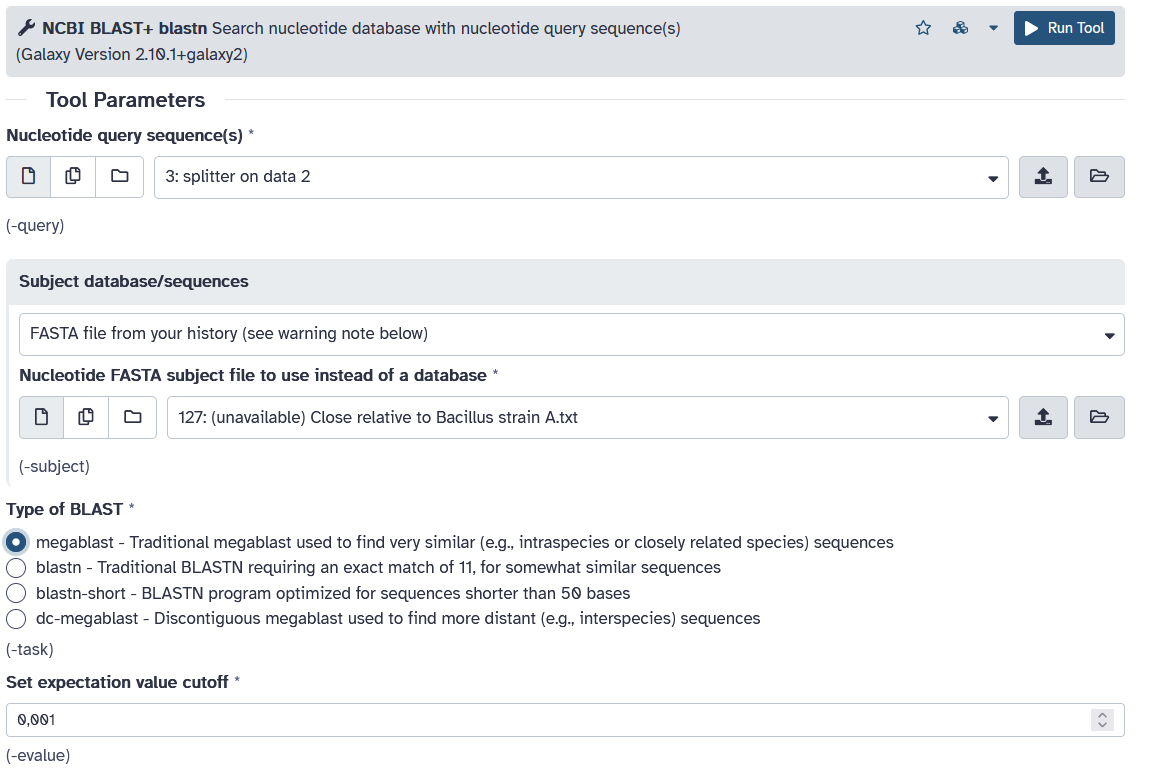

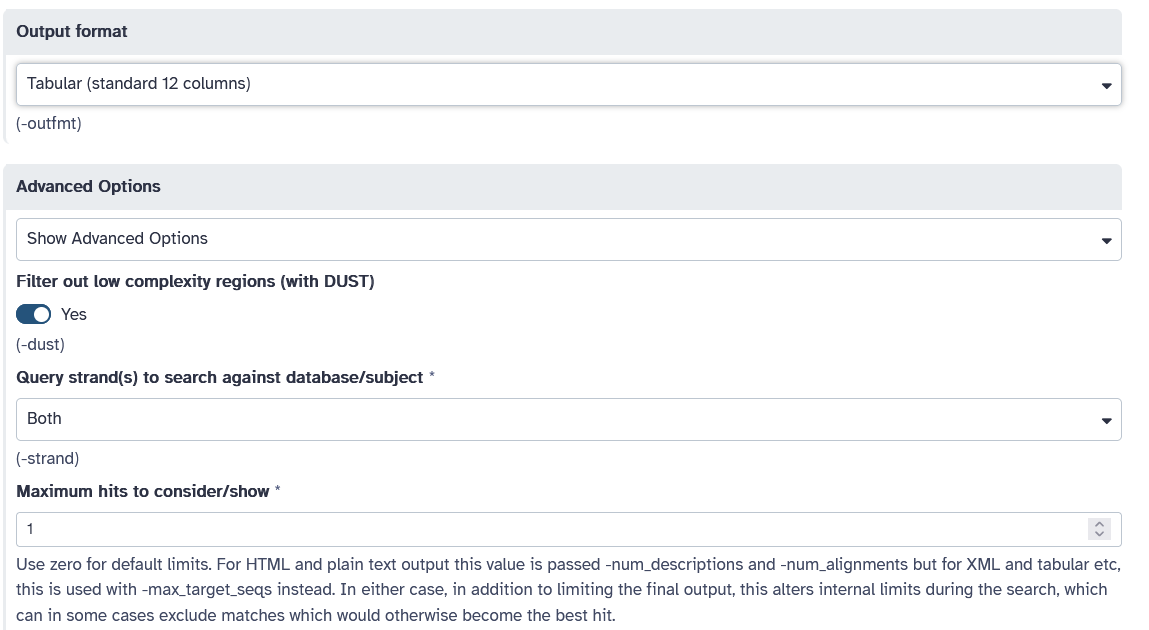

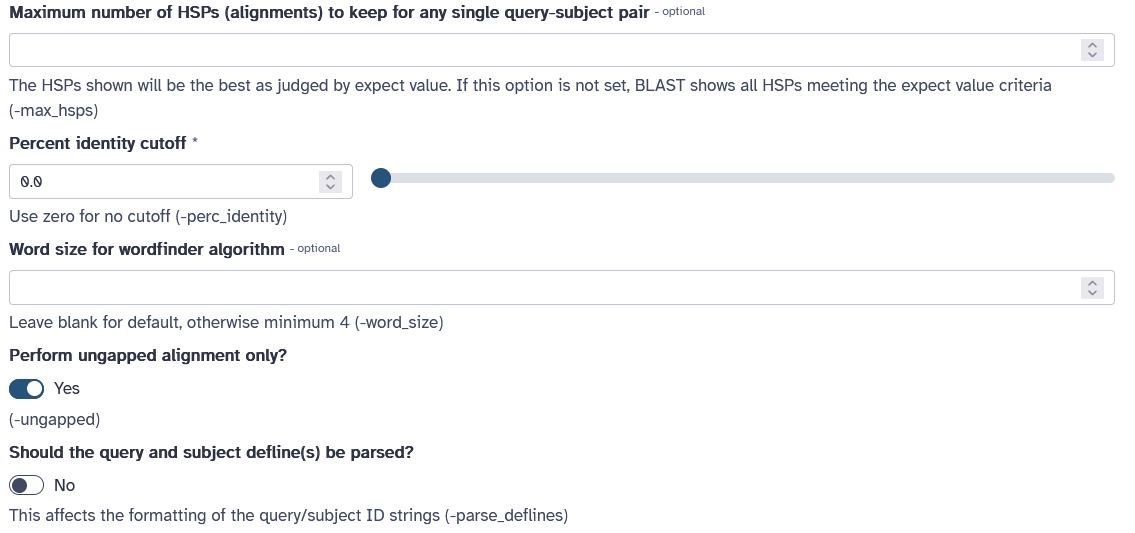

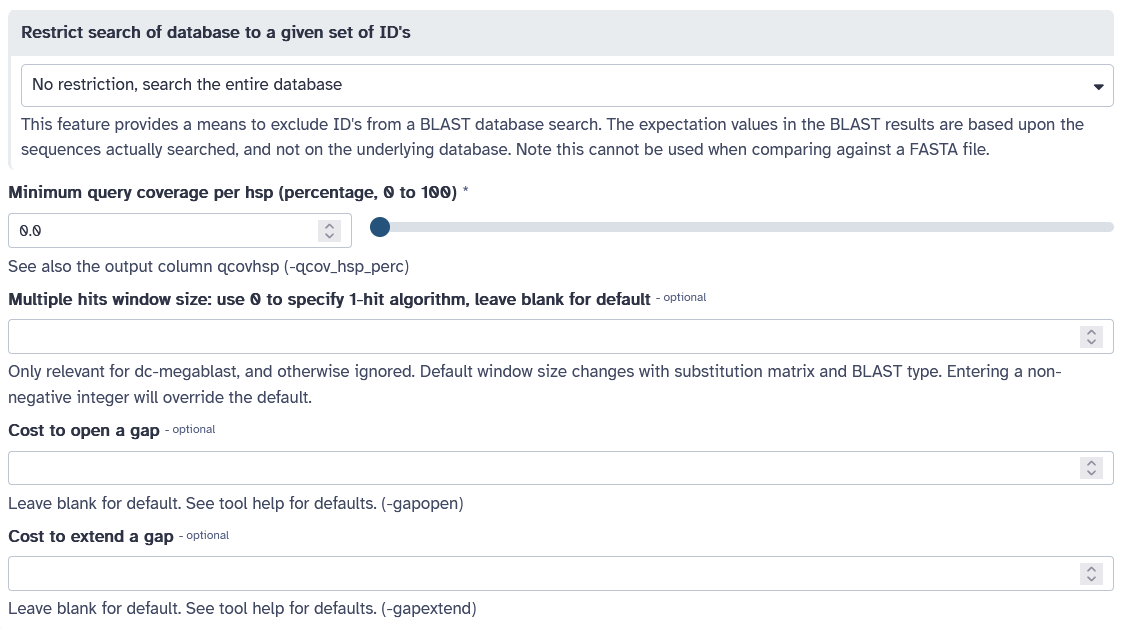

5) Search for the “Compare two datasets” tool and run it on the megablast output and the files with split genome of the strain of interest, using the following parameters:

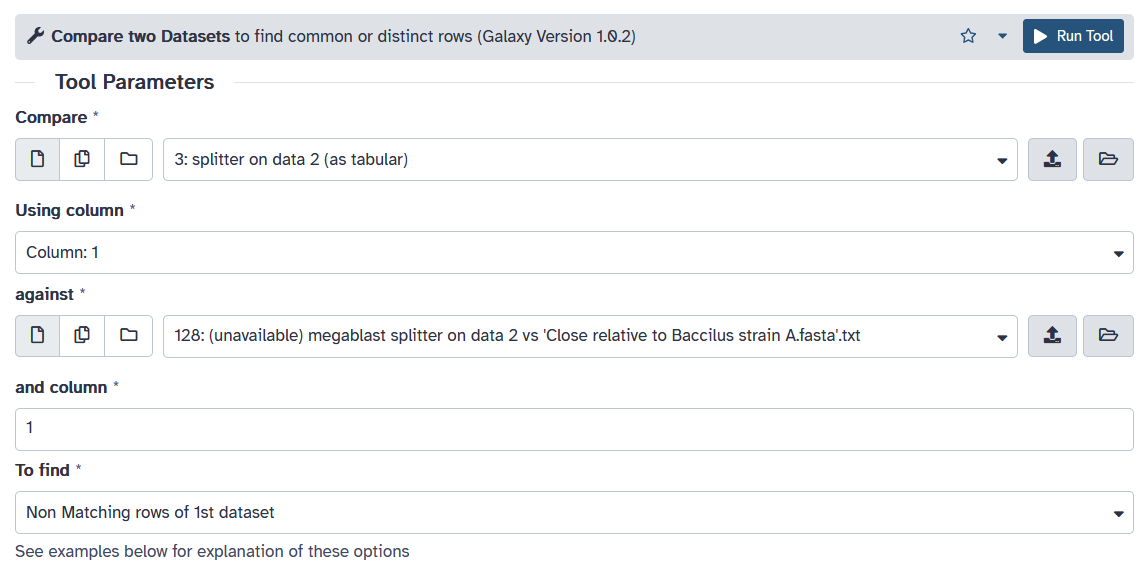

6) Search for the tool “Tabular-to-FASTA” and use it on the output of the previous step using the following parameters:

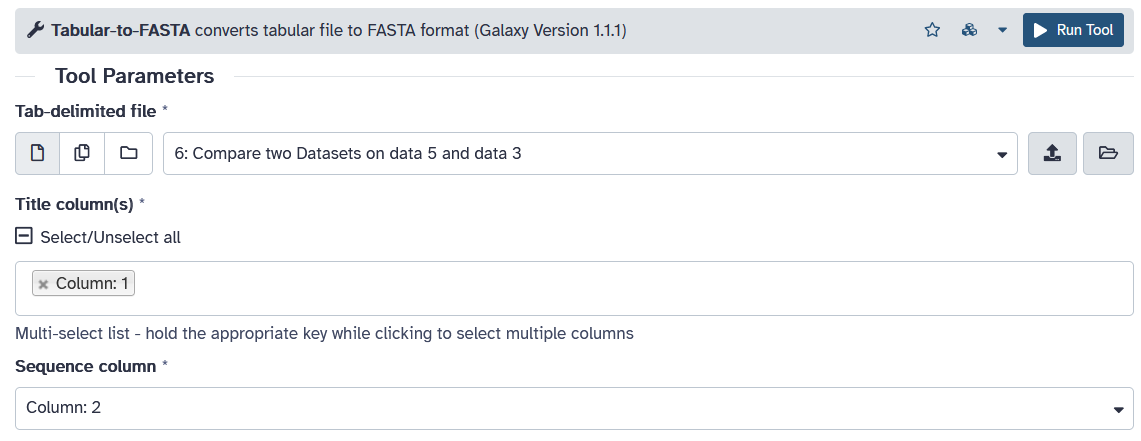

7) Export the files using the “download” button (see red arrow)

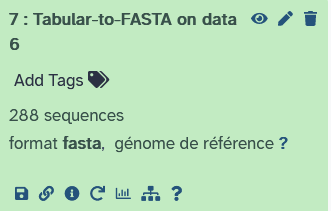

8) Copy and paste the remaining sequences in the nucleotide BLAST tool of NCBI (<https://blast.ncbi.nlm.nih.gov/Blast.cgi>) using the default parameters. If the file has too many sequences, they can be run in smaller batches (it is advised to do it anyway to save time).
9) Discard any sequences with a BLAST hit from the .fasta file.
10) Open the Batchprimer3 tool (https://wheat.pw.usda.gov/demos/BatchPrimer3/), paste the specific sequences and run it with the following parameters:

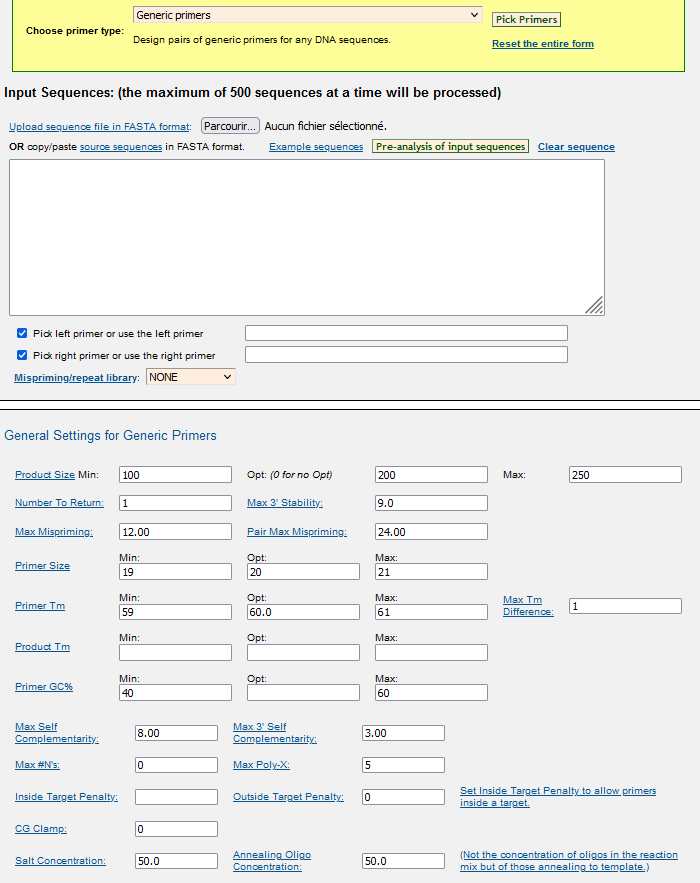

11) Double check the specificity of all the primer pairs using the primer-BLAST tool from NCBI with the following parameters:

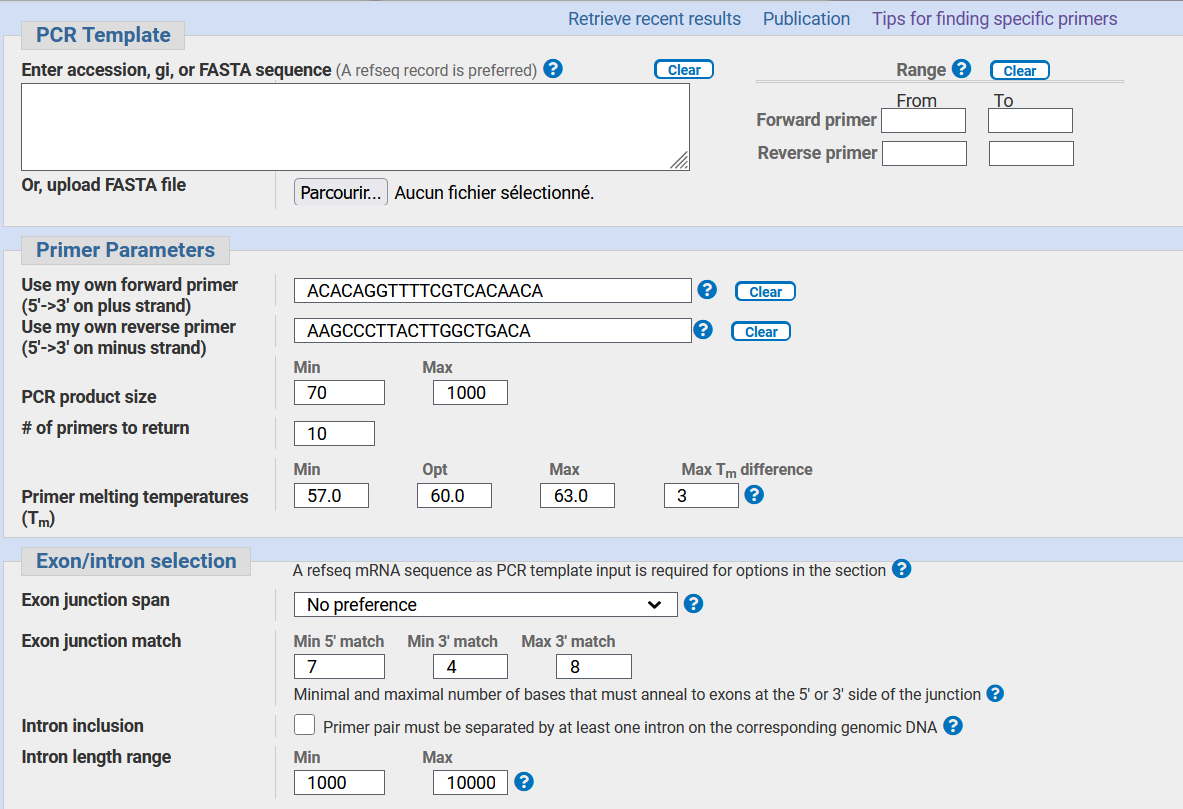

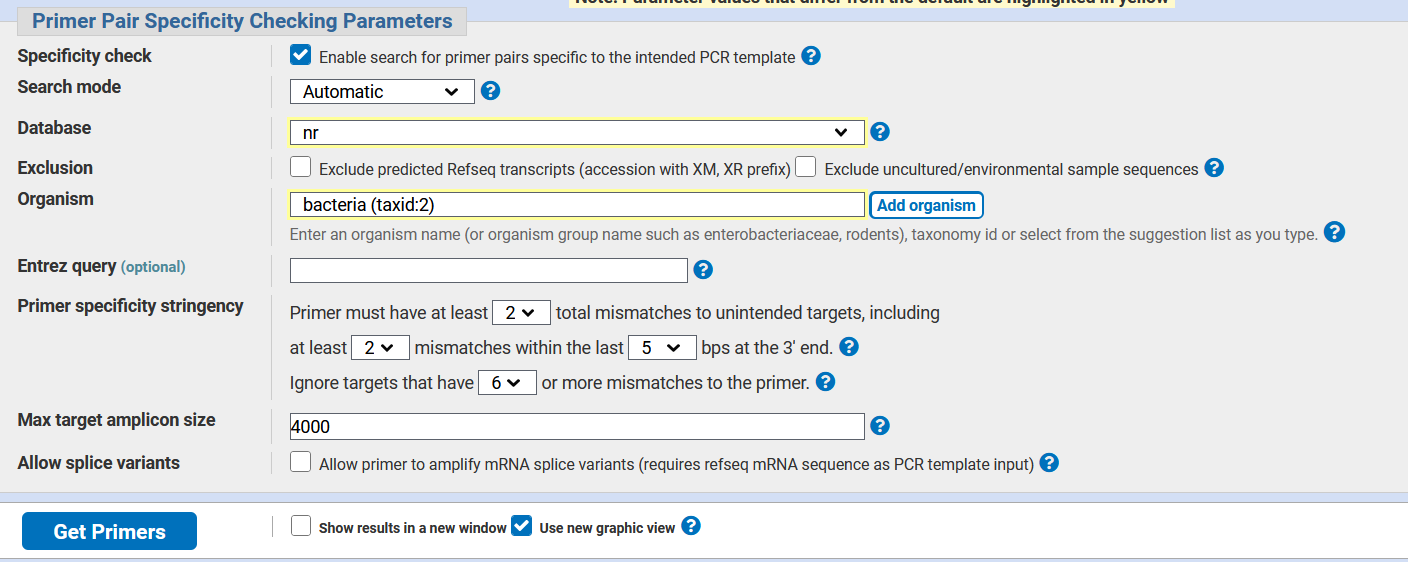

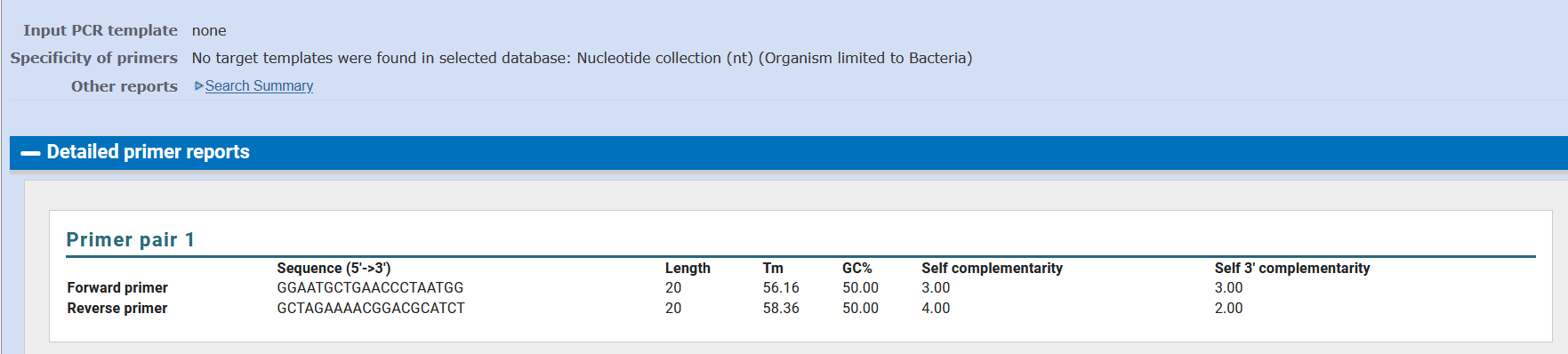

Supplementary file 2: Primers details

| Name | Sequence | Length (bp) | Product size (bp) | GC% | Tm (°C) |
| --- | --- | --- | --- | --- | --- |
| ADPeF1 | CGCTGATCCTCGAAAACTTC | 20 | 199 | 50 | 59.96 |
| ADPeR1 | ATGGGAAGAATGGCTTGATG | 20 |  | 45 | 59.89 |
| 38R_F1 | CCTTAGCGCCAAATAGAACG | 20 | 193 | 50 | 59.87 |
| 38R_R1 | GGATCGATGATGGCTAAACC | 20 |  | 50 | 59.35 |
| TESF1 | GGGTCACTTCTTCCTGGACA | 20 | 196 | 55 | 60.09 |
| TESR1 | CTTGGACTAGGACGCCTTGA | 20 |  | 55 | 60.39 |
| SR38F1 | GTCGTCAACGAGGGTGAGAT | 20 | 198 | 55 | 59.31 |
| SR38R1 | GCAATGACCCAACTCATCAA | 20 |  | 45 | 60.18 |
| AF6 | ACACAGGTTTTCGTCACAACA | 20 | 246 | 50 | 59.5 |
| AR6 | AAGCCCTTACTTGGCTGACA | 20 |  | 50 | 59.4 |
| BF9 | GGAATGCTGAACCCTAATGG | 21 | 119 | 42.8 | 59.1 |
| BR9 | GCTAGAAAACGGACGCATCT | 20 |  | 50 | 59.9 |

Supplementary file 3: PCR analyses of various environmental samples as well as atrazine degrading strain genomic DNA using our specific primers. The figure shows two 2% agarose gels post electrophoresis. On each gel, positive controls (C+) are the respective strain’s genomic DNA. For the PCR, reactions were initiated for 3 min at 95°C, then 35 cycles of the following steps were done: 30 sec at 95°C, 45 sec at 60°C or 62°C, 45 sec at 72°C with a final elongation of 10 min at 72°C.

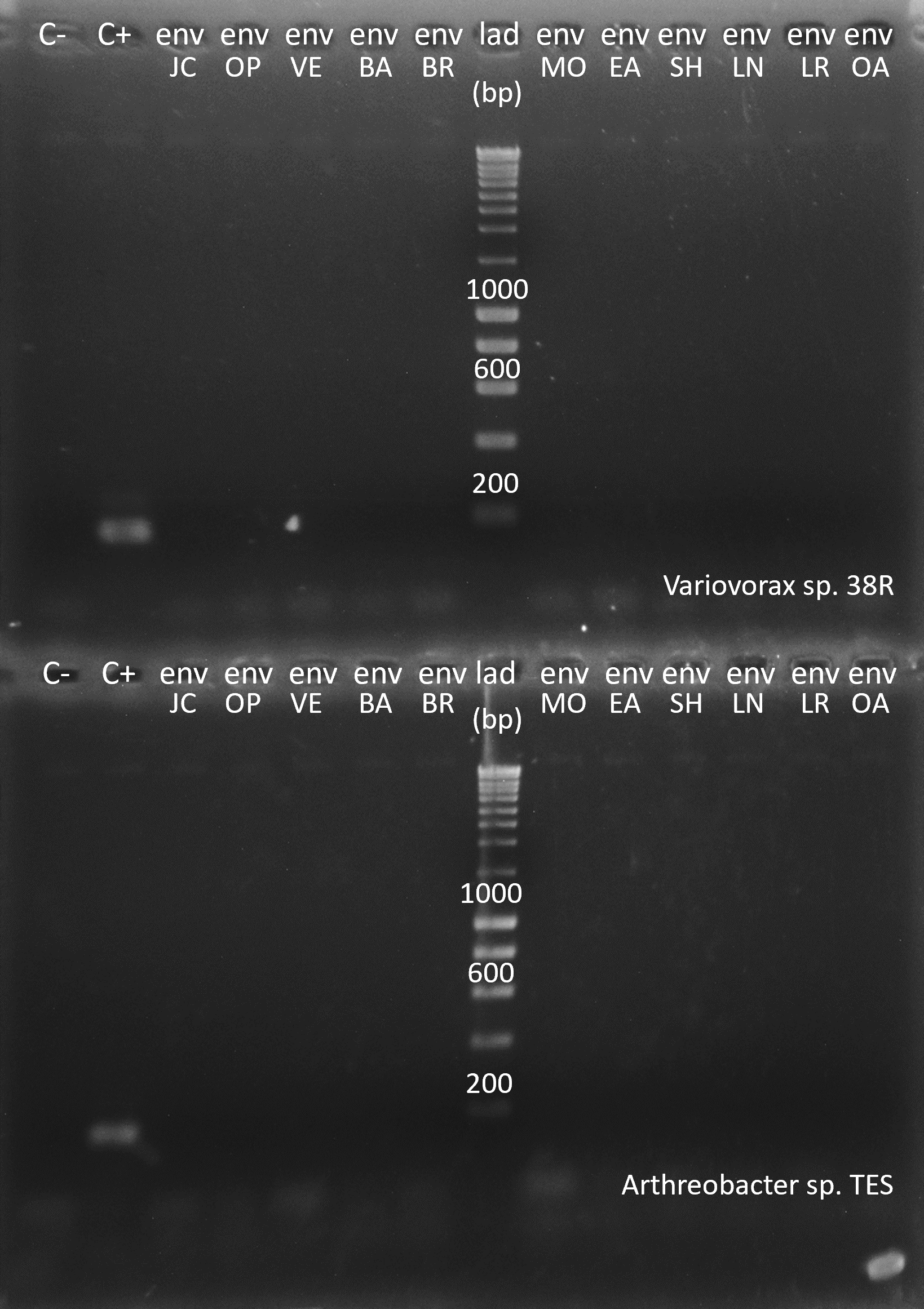

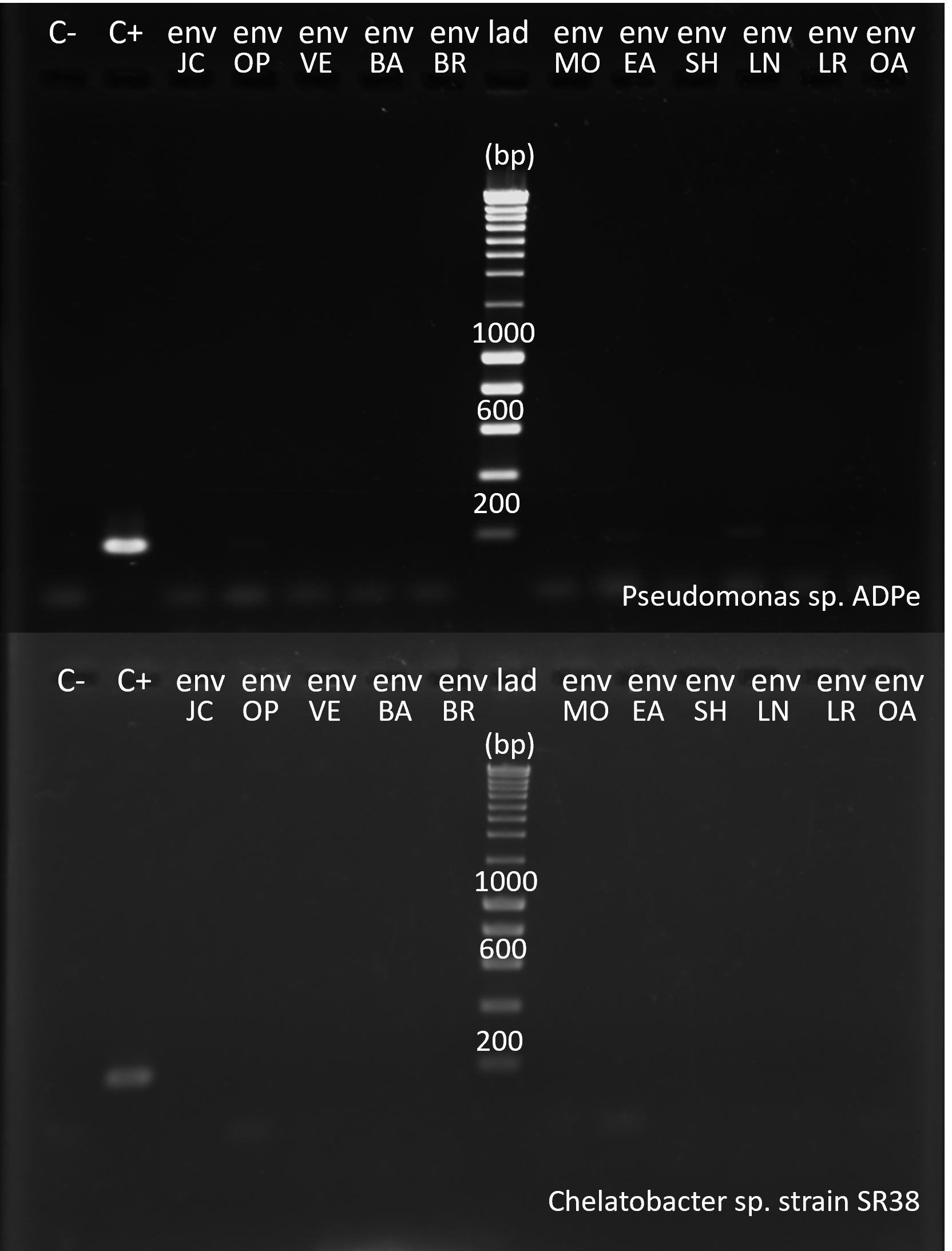

Supplementary file 4: PCR analyses of various environmental samples as well as strain A and B genomic DNA using our specific primers. The figure shows two 2% agarose gels post electrophoresis. To produce the gel on the left, the primer pair AF6/AR6 was used to perform a PCR on environmental DNA samples coming from different locations as well as on the strains genomic DNA. For the right gel, the same analysis was done but using the primer pair BF9/BR9. For the PCR, reactions were initiated for 3 min at 95°C, then 35 cycles of the following steps were done: 30 sec at 95°C, 45 sec at 60°C or 62°C, 45 sec at 72°C with a final elongation of 10 min at 72°C.

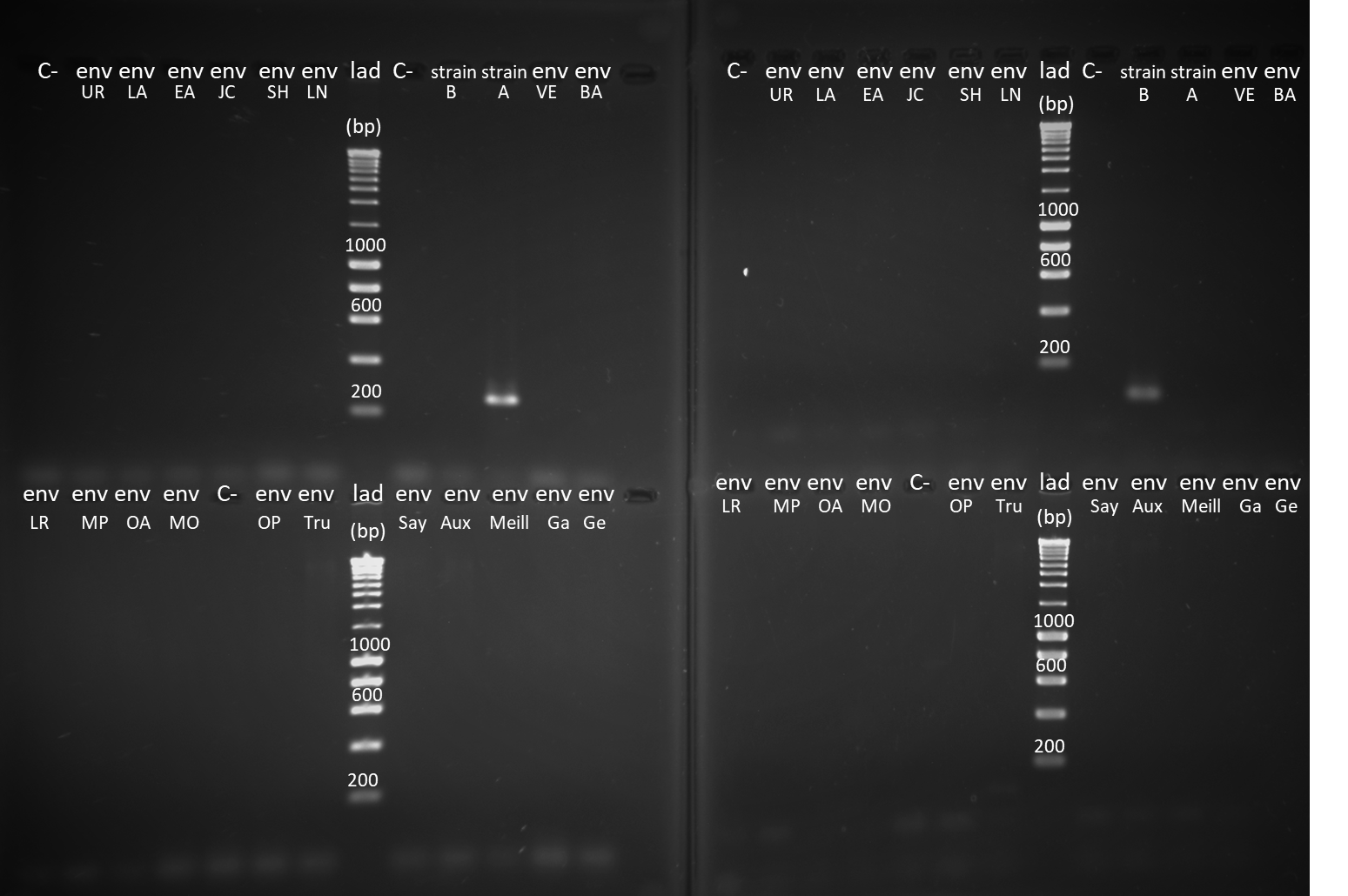

Supplementary file 5: qPCR details

To prepare qPCR standards, the amplicons obtained after running a PCR on genomic DNA using the specific primers AF6/AR6 and BF9/BR9 were cloned in *E. coli* JM109 using pGEMT-easy vector. After the selection of recombinant bacteria, white in colour, a single colony was picked and inoculated in LB medium. Plasmid was purified from an aliquot of the bacterial culture. It was linearized with a restriction enzyme. Its concentration was quantified using a Quantifluor assay to prepare stock standard solutions at 0.5 x 10^9^ copies/µL.

qPCR was performed on a StepOne in 15 µL reactions which comprised 7.5 µL of master mix (Takyon ROX SYBR from Eurogentec),1.5 µL of each primer (10 µM), 2.5 µL of MilliQ water and 2 µL of DNA (0.5 ng/µL). Reactions were initiated for 5 min at 95°C, then 35 cycles of the following steps were done: 15 sec at 95°C, 30 sec at 60°C, 30 sec at 72°C, 20 sec at 72°C for amplification and 15 sec at 95°C, 1 min at 70°C, 15 sec at 95°C for the melt curve.

Supplementary file 6: Standard curves using the specific primers designed on unique sequences of strain A and B

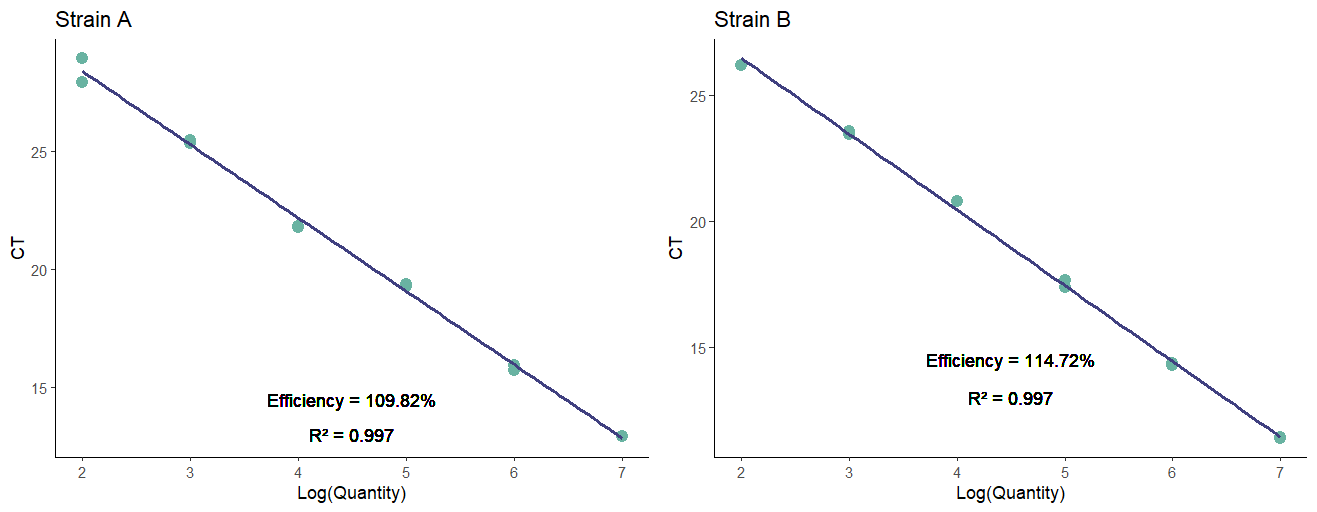

Supplementary file 7: Physico-chemical properties of the soil used for the final validation of the strain A and B specific primers.

| sol name | Clay (%) | Silt (%) | Sand (%) | texture | Ctot (g/kg) | TOC (g/kg) | Ntot (g/kg) | C/N | SOM (g/kg) | pH_water_ |
| --- | --- | --- | --- | --- | --- | --- | --- | --- | --- | --- |
| Garlede | 19,4 | 67,5 | 13,1 | clay loam | 17,980 | 17,980 | 1,476 | 12,200 | 31,100 | 6,070 |
| Gers | 21,1 | 61,3 | 17,6 | clay loam | 36,796 | 36,796 | 2,556 | 14,400 | 63,700 | 5,890 |
| Meillans | 8,1 | 19,3 | 72,6 | sandy | 19,223 | 19,223 | 1,215 | 15,800 | 33,300 | 6,160 |
